# Inheritance of Rootstock Effects in Avocado (*Persea americana* Mill.) cv. Hass

**DOI:** 10.1101/2020.08.21.261883

**Authors:** Paula H. Reyes-Herrera, Laura Muñoz-Baena, Valeria Velásquez-Zapata, Laura Patiño, Oscar A. Delgado-Paz, Cipriano A. Díaz-Diez, Alejandro A. Navas-Arboleda, Andrés J. Cortés

## Abstract

Grafting is typically utilized to merge adapted seedling rootstocks with highly productive clonal scions. This process implies the interaction of multiple genomes to produce a unique tree phenotype. Yet, the interconnection of both genotypes obscures individual contributions to phenotypic variation (*i*.*e*. rootstock-mediated heritability), hampering tree breeding. Therefore, our goal was to quantify the inheritance of seedling rootstock effects on scion traits using avocado (*Persea americana* Mill.) cv. Hass as model fruit tree. We characterized 240 rootstocks from 8 avocado cv. Hass orchards in three regions of the province of Antioquia, in the northwest Andes of Colombia, using 13 microsatellite markers (simple sequence repeats – SSRs). Parallel to this, we recorded 20 phenotypic traits (including morphological, eco-physiological, and fruit yield and quality traits) in the scions for three years (2015–2017). Relatedness among rootstocks was inferred through the genetic markers and inputted in a ‘genetic prediction’ model in order to calculate narrow-sense heritabilities (*h*^*2*^) on scion traits. We used three different randomization tests to highlight traits with consistently significant heritability estimates. This strategy allowed us to capture five traits with significant heritability values that ranged from 0.33 to 0.45 and model fits (*R*^*2*^) that oscillated between 0.58 and 0.74 across orchards. The results showed significance in the rootstock effects for four complex harvest and quality traits (*i*.*e*. total number of fruits, number of fruits with exportation quality, and number of fruits discarded because of low weight or thrips damage), while the only morphological trait that had a significant heritability value was overall trunk height (an emergent property of the rootstock-scion interaction). These findings suggest the inheritance of rootstock effects, beyond root phenotype, on a surprisingly wide spectrum of scion traits in ‘Hass’ avocado. They also reinforce the utility of SSR markers for relatedness reconstruction and genetic prediction of complex traits. This research is, up to date, the most cohesive evidence of narrow-sense inheritance of rootstock effects in a tropical fruit tree crop. Ultimately, our work reinforces the importance of considering the rootstock-scion interaction to broaden the genetic basis of fruit tree breeding programs, while enhancing our understanding of the consequences of grafting.

## INTRODUCTION

How different genomes interact to shape a unique phenotype has been one of the most pervasive questions in quantitative genetics and molecular evolution (Lynch, 2007). Horizontal gene transfer (Bennetzen, 1996) and allopolyploidy (Abbott *et al*., 2013) are often regarded as the typical processes that lead to the interaction of various genomes within a single organism. However, a commonly disregarded yet ancient process that also produces genetic chimeras is grafting, which refers to the agricultural practice that joins the root system (rootstock) of one plant, usually a woody crop, to the shoot (scion) of another (Warschefsky *et al*., 2016; Gautier *et al*., 2019). Grafting started with the earliest tree crops (*i*.*e*. olive, grape, and fig) and rapidly expanded to several Rosaceae (*i*.*e*. apple, plum, pear, and cherry). Nowadays grafting is essential not only for the clonal propagation of highly profitable fruit trees (*i*.*e*. citrus and avocado) but also for the establishment of seed orchards for the wood industry (*i*.*e*. pines, teak). Since grafting is a common practice across a phylogenetically diverse array of fruit and forest trees species, it sets a unique experimental playground to explore the rootstock-scion interaction and enrich our knowledge of chimeric organisms.

Grafting is typically utilized to merge resilient rootstocks to clonal scions that produce the harvested product, either fruits or wood. This way, grafting side steps the bottlenecks of breeding woody perennials, primarily associated with their outcrossing reproductive system and prolonged juvenile phases (Warschefsky *et al*., 2016). The root phenotype may confer direct resilience to root pest and pathogens as well as to abiotic stresses such as drought, flooding and salt soil conditions (Gautier *et al*., 2019). The rootstock can also induce less trivial scion morphological traits such as dwarfing and precocity, and even alter its productivity traits like flowering, fruit set, fruit weight, wood density and pulp yield. Rootstock effects can go further and influence properties typically attributed to the clonal scion such as fruit quality (*e*.*g*. texture, sugar and nutrient content, acidity, pH, flavor, and color), cold tolerance and shoot pest and pathogen resistance (Goldschmidt, 2014). These combined effects are mainly due to large-scale movement of water, nutrients, hormones, proteins, mRNAs and small RNAs (sRNAs) (Wang *et al*., 2017). Despite shared physiological processes account for the overall trait variation, the interconnection of all contributing variables (*i*.*e*. rootstock genotype, scion genotype, and environment) obscures individual contributions to phenotypic variation (Albacete *et al*., 2015; Warschefsky *et al*., 2016). Therefore, an explicit estimation of rootstock effects (*i*.*e*. rootstock-mediated heritability) would be a major advance to speed-up tree breeding programs and discern the consequences of grafting. Narrow-sense heritability (*h*^*2*^), or the proportion of phenotypic variance among individuals in a population due to genetic effects, is regarded as a base-line of any breeding program (Holland *et al*., 2003) since it ensures that genetic gains are maximized per unit time by optimizing breeding and selection cycles (Dieters *et al*., 1995). Yet, in grafted tree species heritability estimation has been hampered by the complexity of the rootstock-scion interaction. A modern marker-based approach to estimate heritability on populations of mixed ancestry is the so-called ‘genetic prediction’ model, which relies on a linear predictor to estimate the additive genetic contribution to phenotypic trait variation and thus trait heritability (Meuwissen *et al*., 2001; Crossa *et al*., 2017). Here we expanded the ‘genetic prediction’ model to a grafted clonal fruit tree by genotyping rootstocks and phenotyping traits at the tree level.

An important fruit tree crop that is nowadays seeing an unprecedented expansion in tropical and subtropical areas is avocado (*Persea americana* Mill.) cv. Hass. Avocado originated in Central America from where it expanded southwards to the northwest Andes, leading to three horticultural races, mid-altitude highland Mexican (*P*. *americana* var. *drymifolia* Schlecht. et Cham. Blake) and Guatemalan (*P*. *american*a var. *guatemalensis* L. Wms.) races, and lowland West Indian (*P*. *americana* var. *americana* Mill.) race (Bergh and Ellstrand, 1986). Previous research about the effect of the selected avocado rootstocks over crop performance has shown that trees of the same variety grafted to Mexican or Guatemalan race rootstocks differ in their susceptibility to *Phytophtora cinnamomi* (Smith *et al*., 2011; Reeksting *et al*., 2016; Sánchez-González *et al*., 2019), in their mineral nutrient uptake (Bard and Wolstenholme, 1997; Calderón-Vázquez *et al*., 2013), and in their response to salinity (Mickelbart and Arpaia, 2002; Raga *et al*., 2014). For instance, Bernstein *et al*. (2001) demonstrated that even in selected rootstocks chosen by exhibiting excellent fruit production under elevated NaCl-condition, there is a wide range of growth sensitivities that results in growth inhibition or growth stimulation under salt-levels typically found at commercial fields. Furthermore, different race rootstocks change the carbohydrate accumulation profile in trees of the same variety, which is known to drive productivity (Whiley and Wolstenholme, 1990), and can ultimately influence alternate bearing, yield components and nutrition on ‘Hass’ avocado (Mickelbart *et al*., 2007). Rootstocks can even affect postharvest anthracnose development (Willingham *et al*., 2001), as well as the blend of biogenic volatile organic compounds emitted by ‘Hass’ (Ceballos and Rioja, 2019), which could be associated with scion pest attraction. Yet, since rootstock-scion interaction works both ways, different scions can have distinct effects on avocado rootstock traits, such as arbuscular mycorrhizal and root hair development (Shu *et al*., 2017).

Despite several studies have provided evidence of avocado rootstock effects on ‘Hass’ crop performance, the genetic identity and the adaptive potential of the rootstocks that are already planted or are being offered by the nurseries remain a major knowledge gap. Additionally, since many ‘Hass’ avocado orchards are yet to be established worldwide in upcoming years, demand for selected rootstocks is reaching its peak, but explicit rootstock effect estimates are still lacking. Therefore, our goal in this study was to quantify inheritance of rootstock effects on a wide spectrum of ‘Hass’ avocado traits by expanding a ‘genetic prediction’ model to genotyped seedling rootstocks. This effort will throw lights on the consequences of grafting, while enhancing avocado rootstock breeding programs.

## MATERIALS AND METHODS

### Plant material

Avocado cv. Hass production areas in Colombia are widely variable in terms of environmental factors such as altitude, solar radiation, relative humidity, temperature, and precipitation. This variability affects avocado production in terms of agronomic behavior, productivity, yield, and fruit quality. In order to discern environmental drivers from rootstock-mediated heritability, we chose eight commercial orchards of avocado cv. Hass at the Antioquia province that have been in production for at least five years and had comparable management for the exportation market. Orchards spanned three different agro-ecological regions, two in the dairy Northern Andean highland plateau, four in the Eastern Andean highland plateau, and two in the South West coffee region (Fig. 1). At each orchard, we selected six randomly distributed blocks with five trees per block (average spacing 7 x 6 m), for a total of 240 trees grafted on seedling rootstocks (Table S1). Orchards were graphically mapped in R v.3.4.4 (R Core Team) using the *Leaflet* package.

**Figure 1.**
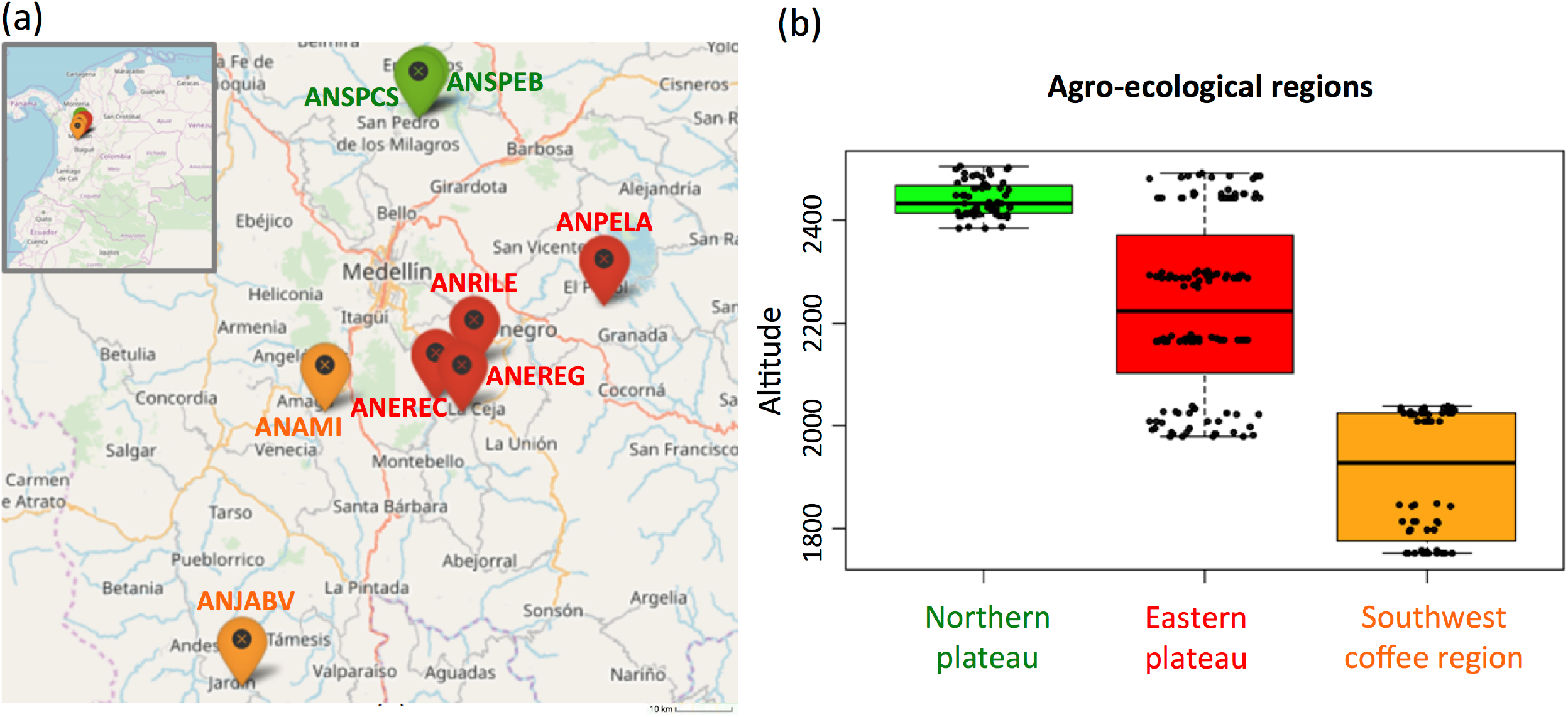
Orchards of ‘Hass’ avocado sampled as part of this study in the northwest Andes of Colombia (province of Antioquia). A total of eight orchards with comparable management for the exportation market spanned three agro-ecological regions, two in the dairy Northern Andean highland plateau (in green), four in the Eastern Andean highland plateau (in red), and two in the South West coffee region (in orange). Thirty trees distributed in six blocks were chosen at each orchard, for a total of 240 trees grafted on seedling rootstocks (Table S1). Orchards names are depicted in (A) while the altitudinal profile per agro-ecological region is shown in (B). The map was done in R v.3.4.4 (R Core Team) using the *Leaflet* package.

### Measurements of phenotypic traits

All 240 trees grafted on seedling rootstocks were measured in 2016 for eight morphological traits. Tree and trunk height were recorded as well as the height of the rootstock and the scion, using the grafting scar as reference. Rootstock and the scion perimeter were measured below and above the grafting scar, too. Trunk perimeter at the grafting scar and a quantitative measure following Webber (1948) were visual proxies for the quality of the grafting. Furthermore, three eco-physiological traits were measured weekly from 2015 to 2017. Flowers and fruits were marked in four cardinally oriented branches, while fallen leaves, flowers and fruits were collected from nets placed aboveground and weighted in order to estimate the total number of leaves, flowers and fruits according to Salazar-García *et al*. (2013). Complete eco-physiological measures were possible for 144 trees across all three years.

Meanwhile, annual harvest from 2015 to 2017 was catalogued in nine categories according to fruit quality. Number of fruits with exportation quality was recorded as a combined trait for yield and quality. If a fruit did not reach quality for exportation, the reason why it was discarded was also annotated. In this sense, the number of fruits that exhibited mechanical or sun damage was recorded, as well as fruits with signs of damage by pests such as scarab beetles (*Astaena pygidialis*) (Holguín and Neita, 2019), thrips (*Frankliniella gardeniae*) or *Monalonion* spp. Furthermore, fruits may not be suitable for exportation due to other imperfections like low weight, early ripening or stalk cut below pedicel, which were annotated, too. Complete harvest categories were possible for 161 trees across all three years.

Trait differences among trees at distinct agro-ecological regions and orchards were determined via Wilcoxon Rank Sum Test for each trait. Additionally, Pearson correlations among phenotypic traits and between them and altitude were calculated using the *PerformanceAnalytics* package. All analyzes were carried out in R v.3.4.4 (R Core Team).

### Genetic screening

Healthy roots from grafted avocado trees were sampled, washed and stored at -20°C. Total genomic DNA was extracted from roots following Cañas-Gutiérrez *et al*. (2015). DNA quality was checked on a Nandrop 2000 (ThermoScientific, U.K). A total of 13 microsatellite markers (simple sequence repeats - SSRs), originally designed by Sharon *et al*. (1997) and Ashworth *et al*. (2004), were chosen for high polymorphism information content (PIC) following estimates by Alcaraz and Hormaza (2007) (Table S2). Forward primers were labeled with WellRed fluorescent dyes on the 5’ end (Proligo, France). SSR markers were multiplexed in three PCR amplifications ran on a Bio-Rad thermo cycler (Bio-Rad Laboratories, Hercules, CA, USA) using the GoTaq® Flexi DNA Polymerase kit (Promega, USA). Reaction volumes and thermocycling profiles were set according to the manufacturer’s instructions. Resulting PCR products were evaluated for thermocycling reaction efficiency on 1.5% agarose gels and then analyzed using capillary electrophoresis in a CEQ 8000 capillary DNA analysis system (Beckman Coulter, Fullerton, CA, USA) at Corporación para Investigaciones Biológicas (CIB, Colombia). Band or alleles sizes were estimated in base pairs with Peak Scanner (Thermo Fisher Scientific, USA) allowing for a maximum of two alleles per sample. High quality genotype data was possible for 188 trees (Table S1), for which DNA extraction, microsatellite amplification and allele scoring succeed.

### Population structure and relatedness estimation

Accuracy of heritability estimates is dependent on population stratification and samples relatedness within populations (Berenos *et al*., 2014; Cortés *et al*., 2014; Sedlacek *et al*., 2016). Therefore, we first assessed population structure with an unsupervised Bayesian clustering approach implemented in STRUCTURE software (Pritchard *et al*., 2000), which determines a *Q* matrix of population admixture across various *K*-values of possible sub-populations found in a sample of genetic diversity more robustly than other cluster methods (Stift *et al*., 2019). A total of 5 independent runs were used for each *K* value from *K*=2 to *K*=7 using an admixture model and 100,000 MCMC replicates with a burn-in of 50,000. Permutations of the output of STRUCTURE were performed with CLUMPP software (Jakobsson and Rosenberg, 2007) using independent runs to obtain a consensus matrix based on 15 simulations. The final structure of the population was determined based on cross-run cluster stability and likelihood of the graph model from Evanno *et al*. (2005), and the admixture index was recorded for each sample.

We further explored within population relatedness using Lynch and Ritland (1999) relatedness estimator because this is the most commonly used, which makes eventual comparisons with other studies easier. Computations were implemented in SPaGeDi v. 1.4 software (Hardy and Vekemans, 2002). Diagonal elements of the matrix were set to one as they describe the relatedness of a genotype with itself. Relatedness estimates between the 28,560 pairwise comparisons were summarized using *hist* and *summary* functions in the R v.3.4.4 (R Core Team) environment.

## Estimation of genetic rootstock effects on scion traits

We used a mixed linear model to predict phenotypic scores for each trait from the rootstock genotypic information following de los Campos *et al*. (2009). Since the scion is clonal, *a priori* scion’s genetic variability was neutral, making possible to highlight the effect of rootstock genetics into the scion phenotype. Thus, we used the additive model described in the equation (1) to predict the phenotypic value based on the rootstock’s genotype.

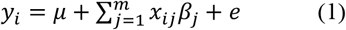

where *yi* is the score predicted for each trait for the *i*^*th*^ individual, *μ* is the mean of each trait in the entire population, *x*_*ij*_ is the relatedness between the *i*^*th*^ and the *j*^*th*^ individuals, following Lynch and Ritland (1999) and Cros *et al*. (2015), *m* is the total number of samples, *β*_j_ is the estimated effect for the relatedness to the *j*^*th*^ individual on the trait and *e* is the estimated error associated with the trait. By using Lynch and Ritland (1999)’s relatedness estimate within equation (1) we are able to enlarge the set of variables to 240. Yet, we still considered a simpler model using the genetic markers by themselves instead of the relatedness matrix, so that *x*_*ij*_ was the genotype of the *i*^*th*^ individual for the *j*^*th*^ marker, and *β*_*j*_ was the estimated maker effect. In order to fit these models to our data we used semi-parametric genomic regression based on reproducing kernel Hilbert spaces regressions (RKHS) methods (Gianola *et al*., 2006; de los Campos *et al*., 2010) implemented in the R package *BGLR* (Perez and de los Campos, 2014). We estimated marker effects and the error associated by running for each trait a Gibbs sampler with 10,000 iterations and an initial burn-in of 5,000.

We computed the narrow sense heritability for each trait following de los Campos *et al*. (2015). We calculated the narrow-sense genetic-estimated heritability h^2^, described in equation (2), as the proportion of phenotypic variance explained by additive effects 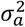 and the sum of 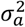 and the random residual 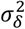. The random residual contains the genetic and environmental effects that cannot be explained by the additive model described in equation (1).

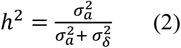

In addition, we estimated the model fit for each trait as the correlation between the trait phenotype and the trait estimation based only on the rootstocks’ relatedness, as shown in equation (3).

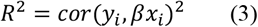

### Permutation tests on phenotype and genotype to obtain significance scores

We used three different permutation tests to obtain significance scores to validate whether scion traits were affected by rootstocks’ genotypes. We permuted the three separate inputs: (1) the observed phenotypic vector – *yi* as in equation (1), (2) the matrix of molecular markers genotyped in the rootstocks – *x*_*ij*_ or the genotype of the *i*^*th*^ individual for the *j*^*th*^ marker as in equation (1), and (3) the matrix of genetic relatedness among rootstocks – *x*_*ij*_ or the relatedness between the *i*^*th*^ and the *j*^*th*^ individuals as in equation (1). In all cases, we used 50 random permutations without replacement, so that the resampling would approximate a random sample (‘null’ distribution) from the original population. We obtained one-sided *p*-values (type I error) for each permutation type, expressed as the proportion of sampled permutations where resultant heritability was larger than the observed heritability estimate. All labels were exchangeable under the null hypothesis. We used this strategy to highlight traits significantly linked with the rootstocks’ genotypes, that is those for which significant *p*-values (*p* < 0.05) were obtained simultaneously for all three types of permutations.

## RESULTS

### Phenotypic differences among agro-ecological regions and orchards

There were significant differences in the distributions of 15 out of 20 phenotypic traits among different agro-ecological regions, according to Wilcoxon Rank Sum Test (*p* < 0.05, Table S3). In general, traits recorded at trees in the South West coffee region had a distribution shifted to the right compared to trees in the Northern and Eastern Andean highland plateaus. The three measures of perimeter (in the rootstock, scion and trunk) and the number of fruits with mechanical damage from trees in the Northern plateau had a median higher than trees in the South West coffee region and the Eastern plateau. Trees in the South West region exhibited higher medians for four out of eight morphological traits (tree height, trunk height, rootstock length, and rootstock compatibility), two out of three eco-physiological measures (number of fruits and number of leaves), and five out of nine annual harvest traits (number of fruits with exportation quality, low weight and sun damage, as well as those damaged by thrips or ripened) (*p* < 0.05, Table S3).

Meanwhile, there were differences in the distributions of seven, nine, and 19 traits between orchards within the Northern, South West and Eastern agro-ecological regions, respectively, based on Wilcoxon Rank Test (*p* < 0.05, Table S3). Orchards with highest trait’s medians were ANSPEB and ANPELA in the Northern and Eastern plateaus, respectively. Details regarding trait’s distribution differences by regions and orchards are depicted in Fig. S1-S5.

Regarding altitude, there were significant trait differences for 14 out of 20 traits (*p* < 0.05, Table S5). For all cases, the correlation with the altitude was negative. The strongest altitudinal correlations were for the rootstock (*R*^*2*^ = -0.61, *p* < 0.05) and trunk (*R*^*2*^ = -0.58, *p* < 0.05) heights, and the number of fruits with low weight (*R*^*2*^ = - 0.59, *p* < 0.05).

Finally, most of these traits were also significantly correlated with each other. In the group of morphological traits, the highest correlations were between (1) tree height and scion length (*R*^*2*^ = 0.94, *p* < 0.05, Fig. S6), and (2) the perimeters of the rootstock, scion and the overall trunk (*R*^*2*^ = 0.8 – 0.84, *p* < 0.05, Fig. S6). The three eco-physiological traits had medium correlations (*R*^*2*^ = 0.37 – 0.40, *p* < 0.05, Fig. S7). For harvest traits, the highest correlations were between (1) the number of fruits with the stalk cut below the pedicel and with damage caused by thrips (*R*^*2*^ = 0.64, *p* < 0.05, Fig. S8), and (2) the number of fruits with low weight and with exportation quality (*R*^*2*^ = 0.61, *p* < 0.05, Fig. S8).

### Relatedness and population structure estimates

Evaluation of population structure using an unsupervised Bayesian clustering approach implemented in STRUCTURE with *K*=2 to *K*=10 sub-populations resulted in an ideal *K*-value of 3 sub-populations (Fig. S9) based on the increases in likelihood ratios between runs using Evanno’s delta *K* statistic (Evanno *et al*., 2005) and cross-run cluster stability. Points of inflection were not observed for the log-likelihood curve but a smaller increase of the likelihood was found when comparing *K*=3 and *K*=4 to other *K*-values. Yet, cross-run cluster stability did not result in the split of a fourth sub-population compared to *K*=3.

Separation of the sub-populations at each *K*-value is informative and therefore is presented in Fig. 2. At the first level of sub-population separation, *K*=2, one orchard from the Northern plateau (ANSPEB) split, while the other orchard from the Northern plateau (ANSPCS) and two from the Eastern plateau (ANEREC and ANEREG) revealed high levels of admixture. At *K*=3 two orchards from the Eastern plateau (ANEREC and ANPELA) differentiated from the others by high levels of admixture. At *K*=4 all sub-populations were admixed for the fourth sub-population but ANSPEB, which differentiated homogeneously since *K*=2. Higher K-values did not contribute further divergence but increased overall admixture levels.

**Figure 2.**
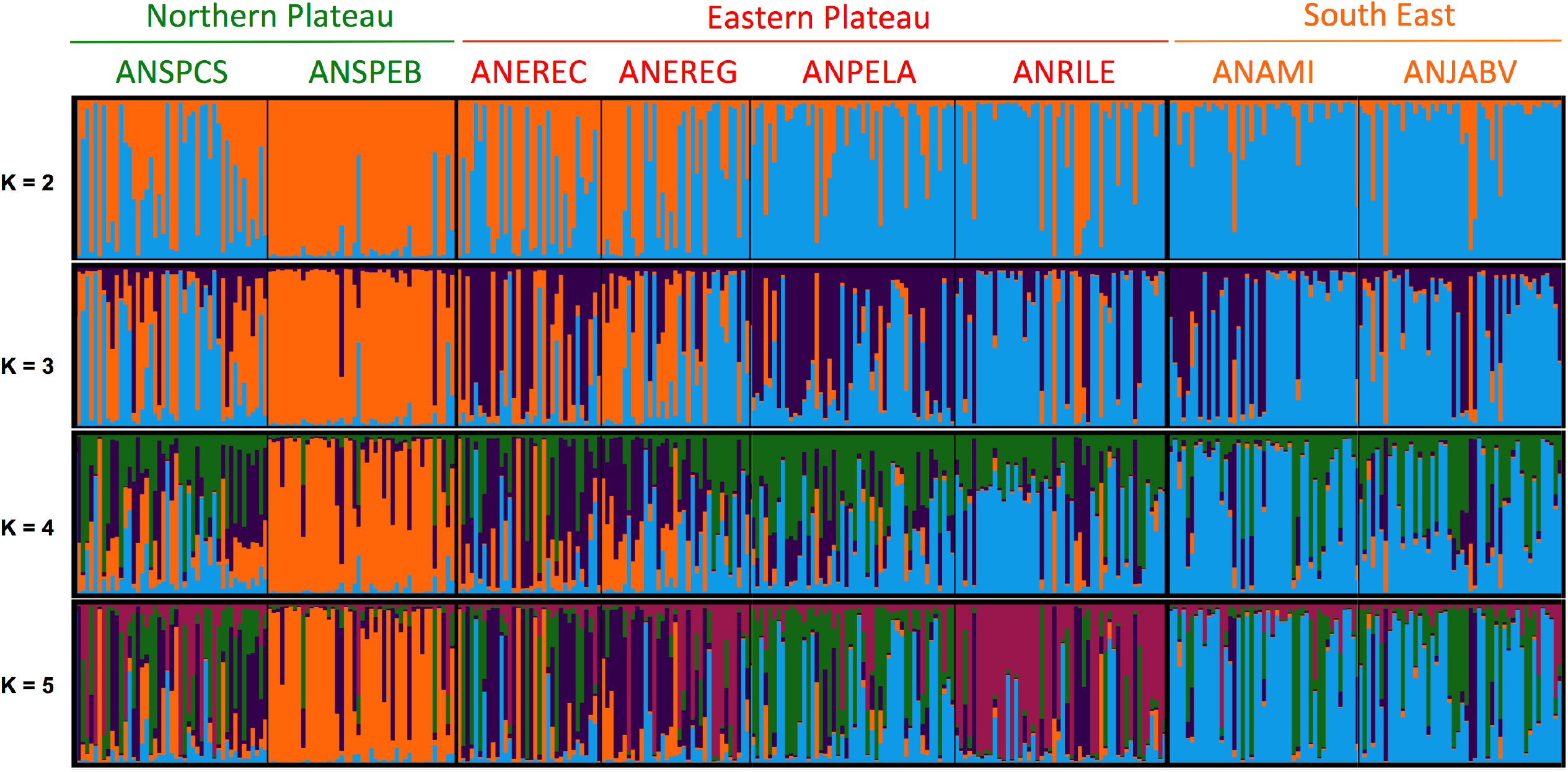
Population structure of seedling rootstocks across eight orchards of ‘Hass’ avocado as inferred with an unsupervised Bayesian clustering approach implemented in STRUCTURE (Pritchard *et al*., 2000) using 13 SSR markers. Orchards are sorted and colored according to the agro-ecological region, and their names are shown at the top of the bar plot. *K*-values of possible sub-populations ranged from 2 to 5. The optimum *K*-value of 3 was determined based on cross-run cluster stability of five independent runs and likelihood of the graph model (Fig. S9) from Evanno *et al*. (2005). Higher *K*-values did not contribute further divergence yet increased overall admixture levels. The *Q* matrix of population admixture at *K* = 3 and the admixture levels are summarized in Table S6. The SSR markers (2) were designed by Sharon *et al*. (1997) and Ashworth *et al*. (2004), and were prioritized according to their polymorphism information content (PIC) following Alcaraz and Hormaza (2007).

Admixture levels at *K*=3 in the orchard of the Northern plateau (ANSPCS) and the two orchards of the Eastern plateau (ANEREC, ANEREG) that exhibited high heterogeneity from *K* = 2 were significantly higher than in the rest (0.26 ± 0.05 *vs*. 0.18 ± 0.03, *p* < 0.05, Table S6). The more distant orchard (ANSPEB) was the less admixed (0.10 ± 0.03). Overall non-zero genetic distance according to Lynch and Ritland (1999) ranged from 0.2 to 1.0 (Fig. S10).

### Genetic heritability and predictive ability

Estimates of rootstock-mediated heritabilities (*h*^*2*^) were significant for five of the 20 measured traits (Fig. 3), regardless the permutation strategy (Fig. 4), and ranged from 0.34 to 0.45 averaged *h*^*2*^ values with average model fits (*R*^*2*^) ranging from 0.61 to 0.73 (Table 1). The majority of traits with significant rootstock-mediated heritability were annual harvest traits (number of fruits with exportation quality, low weight, and thrips’ damages with average *h*^*2*^ values of 0.36, 0.35, and 0.34, and average *R*^*2*^ values of 0.61, 0.63 and 0.61, respectively). Only one morphological trait had significant results according to the permutation tests – trunk height with average *h*^*2*^ and *R*^*2*^ values of 0.36 and 0.64. The number of fruits was the only eco-physiological trait that had significant results with average *h*^*2*^ and *R*^*2*^ values of 0.45 and 0.73. In general, significant morphological and physiological traits had higher *h*^*2*^ values (*h*^*2*^ = 0.36 ± 0.01 and *h*^*2*^ = 0.45 ± 0.01 for trunk height and the number of fruits, respectively) than annual harvest traits (*h*^*2*^ = 0.36 ± 0.02, *h*^*2*^ = 0.34 ± 0.01 and *h*^*2*^ = 0.35 ± 0.01 for the number of fruits with exportation quality, low weight and damages caused by thrips, respectively). Meanwhile, trait predictability was high, especially for the significant eco-physiological trait total number of fruits (*R*^*2*^ = 0.73), and was lowest for number of fruits with exportation quality (*R*^*2*^ = 0.58). A marker-based model was statistically unpowered for all traits (Fig. S11).

**Table 1.**
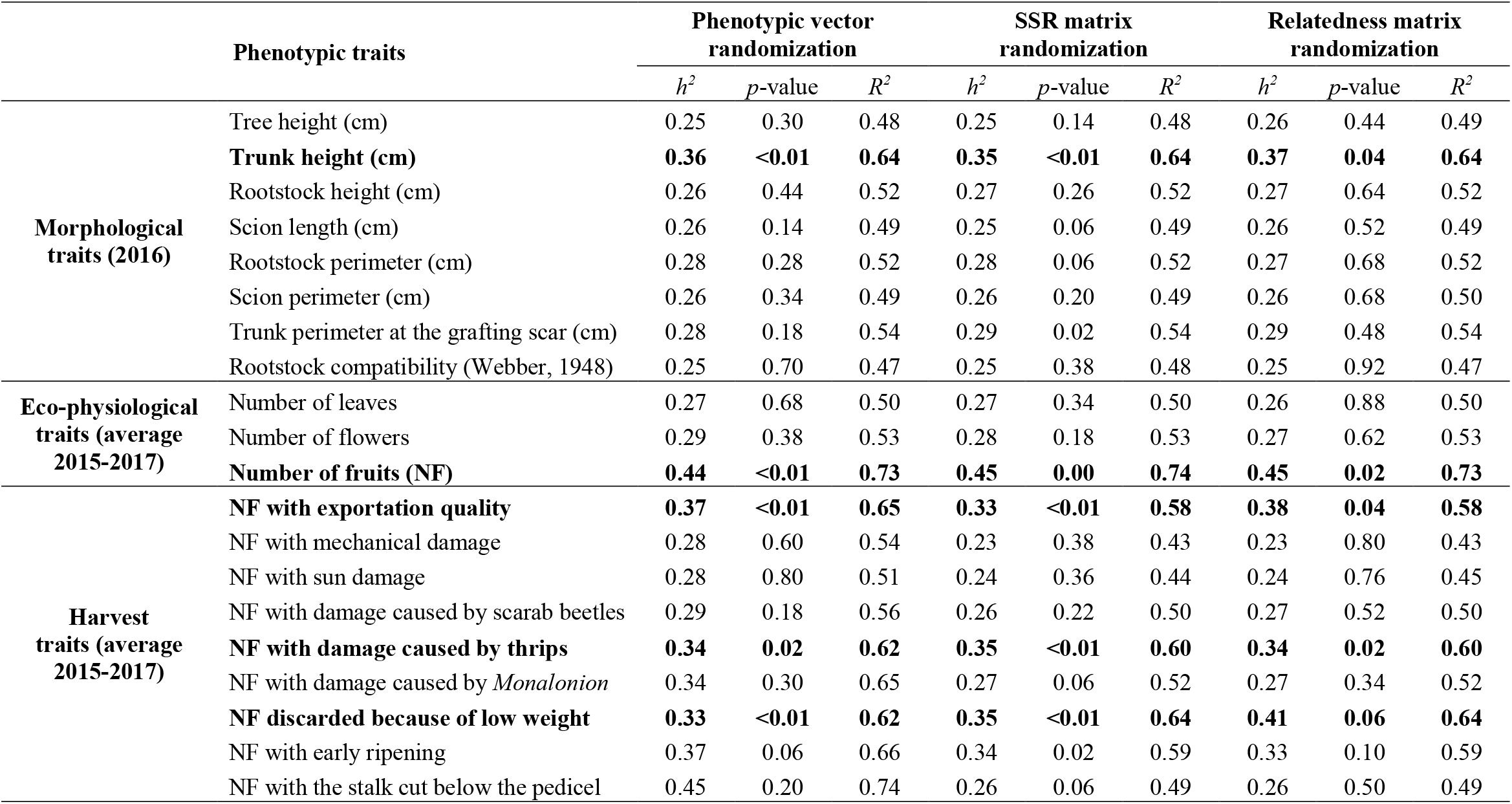
Narrow-sense rootstock-heritability (*h*^*2*^) estimates for the 20 measured traits from eight ‘Hass’ avocado orchards. Heritability (*h*^*2*^) and model fits (*R*^*2*^) estimates were gathered using Lynch and Ritland (1999)’s relatedness matrix inputted in a ‘genetic prediction’ additive mixed linear model, according to de los Campos *et al*. (2009). One-sided *p*-values of the observed heritability were estimated using independent permutations of the phenotypic vector, the matrix of molecular markers and the matrix of genetic relatedness among rootstocks (Fig. 4). Consistently significant values are in bold (Fig. 3).

**Figure 3.**
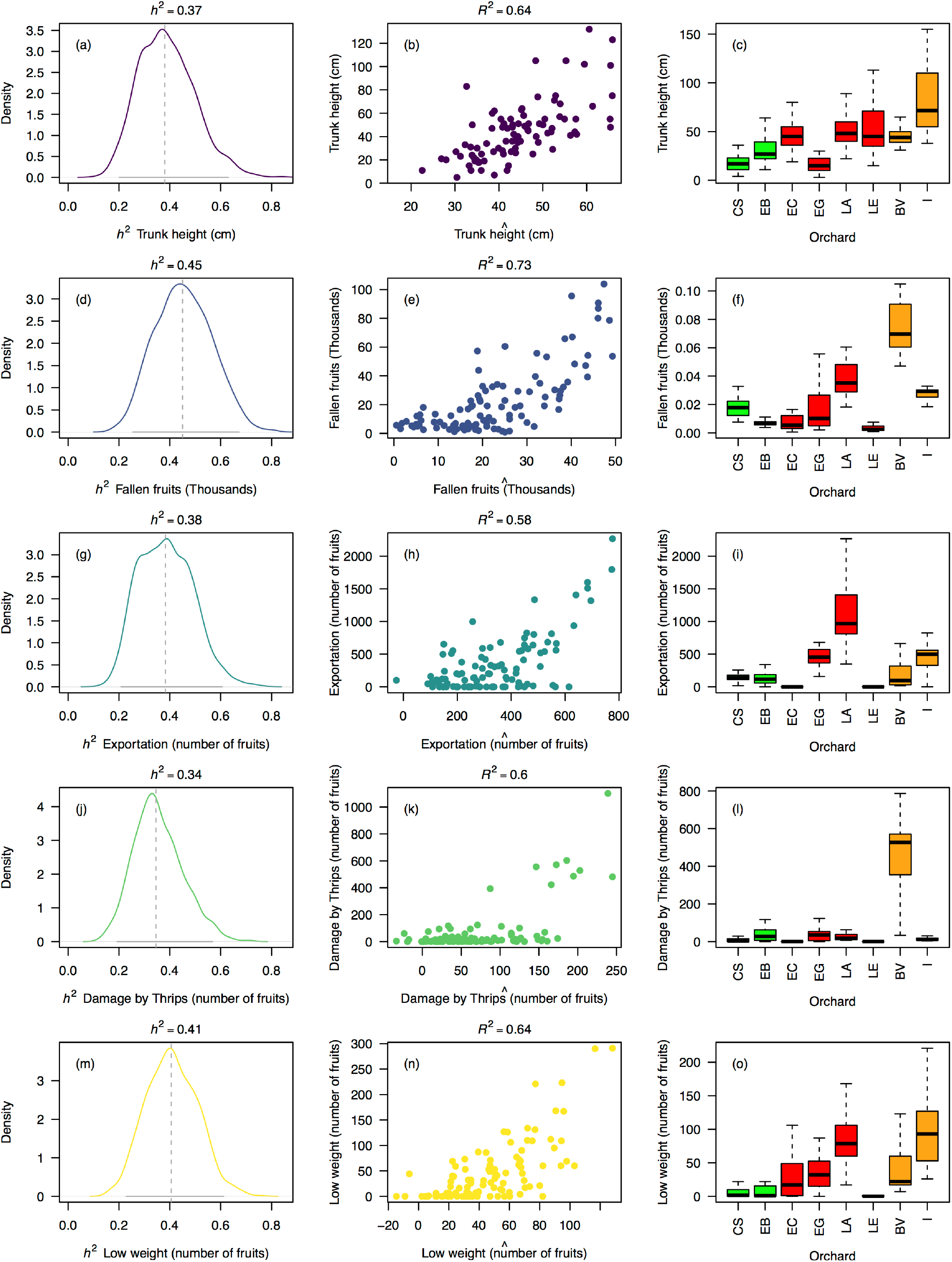
Significant estimates of narrow-sense rootstock-mediated heritability (*h*^*2*^) in five of the 20 measured traits based on a ‘genetic prediction’ model calibrated with Lynch and Ritland (1999)’s relatedness matrix among rootstocks from eight ‘Hass’ avocado orchards. Depicted traits (rows) are those for which significant *p*-values (*p* < 0.05) were simultaneously obtained for three different permutation strategies (of the phenotypic vector, the matrix of molecular markers and the matrix of genetic relatedness among rootstocks, Table 1), although the graphical results only reflect estimates obtained after permuting the relatedness matrix. The first column of figure panels shows the posterior distribution for the rootstock-mediated heritability (*h*^*2*^) estimates as well as their mean (dashed vertical gray line) and 95% confidence interval (continuous horizontal gray line). The second column of figure panels reflects the model fits (*R*^*2*^) expressed as the correlation between the observed trait phenotype (*yi*) and the model’s trait estimation (*βx_i_*) – equation (3). The third column of figure panels recalls the trait distribution across orchards (from Fig. S1-S5). Estimates of *h*^*2*^ and *R*^*2*^ are derived from an additive mixed linear model according to de los Campos *et al*. (2009).

**Figure 4.**
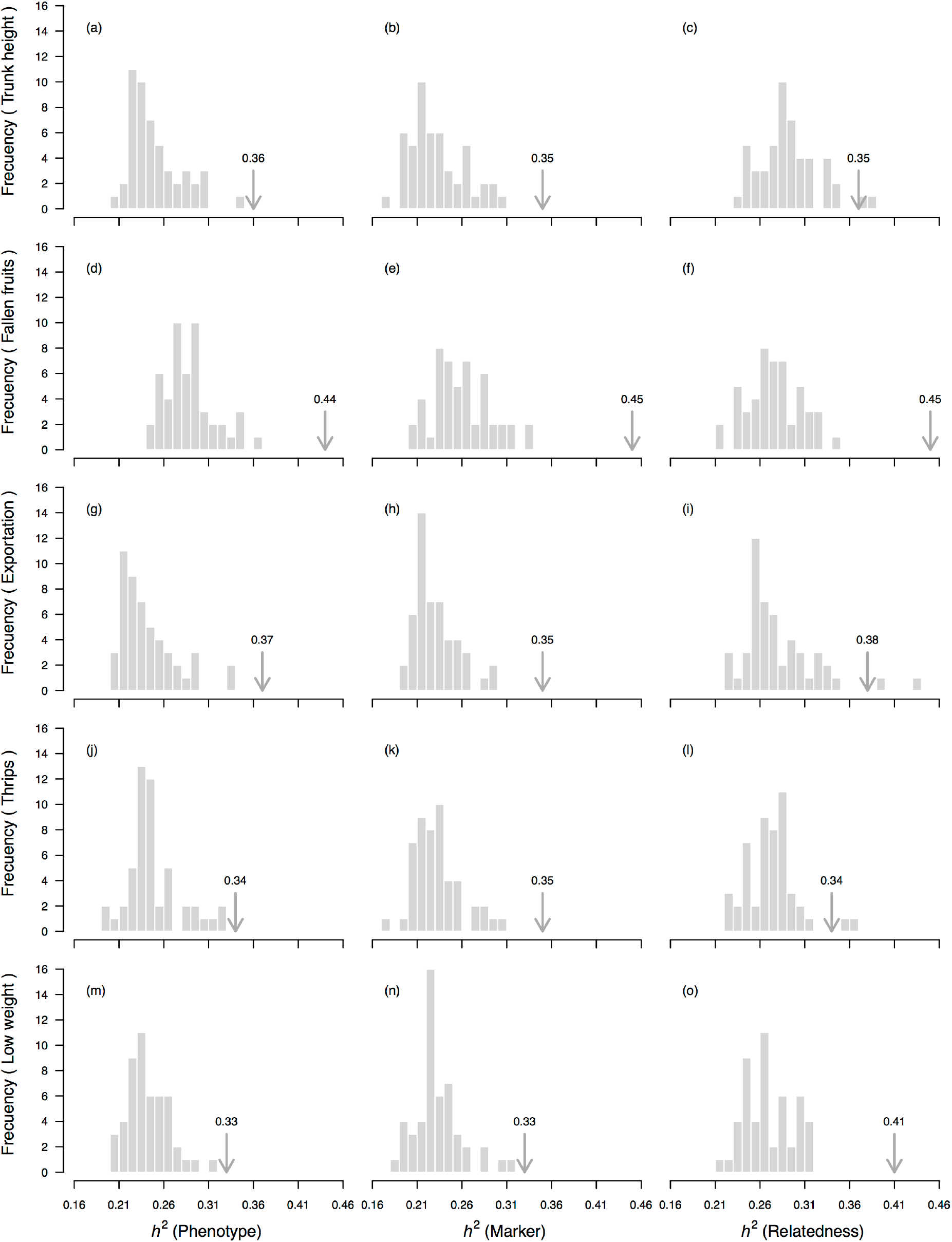
‘Null’ distributions (random sample) of the rootstock–mediated heritability (*h*^*2*^) estimate for five of the 20 measured traits based on a ‘genetic prediction’ model calibrated with Lynch and Ritland (1999)’s relatedness matrix among seedling rootstocks from eight ‘Hass’ avocado orchards. Depicted traits (rows) are those for which significant *p*-values (*p* < 0.05) were simultaneously obtained for three different permutation strategies – of the phenotypic vector (first column of figure panels), the matrix of molecular markers (second column of figure panels) and the matrix of genetic relatedness among rootstocks (third column of figure panels). In all cases 50 random permutations without replacement were used. Average rootstock–mediated heritability (*h*^*2*^) estimates (from Table 1) are marked with an arrow. The proportion of sampled permutations where resultant heritability was larger than the observed heritability estimate corresponds to the one-sided *p*-value reported in Table 1.

## DISCUSSION

We quantified genetic effects of avocado seedling rootstocks on 20 ‘Hass’ scion traits using a ‘genetic prediction’ model that related traits’ variation with the SSR identity of rootstocks from eight different orchards. Trees exhibited high levels of admixture across orchards, consistent with rampant gene flow among putative races. Genetic estimates of rootstock-mediated heritability (*h*^*2*^) were significant for 5 of the 20 measured traits and ranged from 0.33 to 0.45 *h*^*2*^ with model fits (*R*^*2*^) between 0.58 and 0.74 across orchards. The only morphological trait that we found having a significant genetic-estimated heritability value was trunk height, likely an emergent property of the rootstock-scion interaction. Yet, there were significant rootstock effects for various harvest and quality traits such as total number of fruits, number of fruits with exportation quality, and number of fruits discarded because of low weight and thrips damage. These findings suggest the inheritance of rootstock effects on an overwhelming wide spectrum of ‘Hass’ avocado traits relevant for productivity, which will be critical to meet the demands of the growing worldwide market. Our results also reinforce the importance of plant grafting to increase yields, beyond making plants more resistant to disease and abiotic stress in a changing climate.

### Relatedness and population admixture are consistent with rampant gene flow among three populations

Examination of population structure using an unsupervised Bayesian clustering approach and within population relatedness using Lynch and Ritland (1999) relatedness estimator are indicative of three major clusters with high levels of admixture. These clusters likely match the three horticultural races described for avocado, which are mid-altitude highland Guatemalan (*P*. *americana* var. *guatemalensis* L. Wms.) and Mexican (*P*. *americana* var. *drymifolia* Schlecht. et Cham. Blake) races, and lowland West Indian (*P*. *americana* var. *americana* Mill.) race. Genetic analyses and commercial traits have both supported this race structure.

Previous genetic characterizations providing tangential signals of horticultural races have used targeted genes (Chen *et al*., 2009), cpDNA (Ge *et al*., 2019), SSR (Alcaraz and Hormaza, 2007; Ferrer-Pereira *et al*., 2017; Boza *et al*., 2018; Sánchez-González *et al*., 2020) and SNP markers (Kuhn *et al*., 2019c; Rubinstein *et al*., 2019; Talavera *et al*., 2019), in some cases using gene-bank accessions, like from the Venezuelan germplasm bank (INIA-CENIAP) (Ferrer-Pereira *et al*., 2017), the National Germplasm Repository (SHRS ARS USDA) in Miami (Kuhn *et al*., 2019a; Kuhn *et al*., 2019b), and the Spanish germplasm bank (Talavera *et al*., 2019). Despite some of these analyses captured all three races (Talavera *et al*., 2019), others exhibited mixed and inconclusive population structure (Cañas-Gutiérrez *et al*., 2015; Cañas-Gutiérrez *et al*., 2019; Cañas-Gutierrez *et al*., 2019). However, modern genomic tools have not only reinforced race substructure (Rendón-Anaya *et al*., 2019; Talavera *et al*., 2019), but also provided evidence for the hybrid origin of commercially important varieties such as Mexican/Guatemalan ‘Hass’ avocado (Rendón-Anaya *et al*., 2019). Our characterization has further highlighted the admixed origin of seedling rootstocks currently used at commercial orchards in the northwest Andes. Persistent admixture due to rampant gene flow is expected for a species that, as avocado, has been subjected to continent-wide human-mediated migration (Bergh and Ellstrand, 1986; Galindo-Tovar *et al*., 2007), besides being an obligate outcrossing (via protogynous dichogamy, a sequential non- overlapping hermaphroditism in which female function precedes male function).

Regarding economical traits, Guatemalan race typically has small seeds and exhibits late fruit maturity, while Mexican race shows early fruit maturity and cold tolerance. In contrast, West Indian race has large fruit size and low oil content (Bergh and Ellstrand, 1986). However, trait differentiation could not be assessed in this study since genotyping was carried out on seedling rootstocks. In order to evaluate in more detail rootstocks’ fruit phenotype, stooling or layering would need to be induced from rootstocks (Knight *et al*., 1927; Webster, 1995), a technique normally used for clonal propagation of a desired rootstock rather than high-scale phenotyping. A so far unexplored, yet promising alternative, would be to calibrate Genomic Prediction (GP) (Crossa *et al*., 2017) and Machine Learning (ML) (Gianola *et al*., 2011; Libbrecht and Noble, 2015; Schrider and Kern, 2018) models using high-throughput genotyping of phenotyped un-grafted avocado trees spanning all three races, in order to predict rootstocks’ own unobserved phenotypes.

### Significant rootstock effects for various complex harvest and quality traits

Our results suggest the inheritance of rootstock effects on a surprisingly wide spectrum of ‘Hass’ avocado genetically complex traits, mostly spanning economically relevant attributes such as total number of fruits, number of fruits with exportation quality, number of fruits discarded because of low weight, and number of fruits damaged by thrips. The only morphological trait that we found having a significant heritability value mediated by the rootstock was trunk height. Interestingly, all these traits refer to the ability of the rootstocks to impact the phenotype of the grafted scion (*i*.*e*. harvest/quality traits), or the entire tree (*i*.*e*. trunk height), but not the root phenotype itself (*e*.*g*. rootstock height or perimeter). This speaks for a predominant role of the rootstock-scion interaction rather than independent additive effects of each genotype, which is expected when combined effects are mainly due to transport of water and nutrients, and large-scale movement of hormones, proteins, mRNAs and sRNAs (Wang *et al*., 2017).

Previous research about the effect of rootstocks on avocado crop performance has focused on susceptibility to *P*. *cinnamomi* (Smith *et al*., 2011; Reeksting *et al*., 2016; Sánchez-González *et al*., 2019), mineral nutrient uptake (Bard and Wolstenholme, 1997; Calderón-Vázquez *et al*., 2013), and response to salinity (Bernstein *et al*., 2001; Mickelbart and Arpaia, 2002; Raga *et al*., 2014). However, harvest/quality traits have not been explicitly considered in previous studies that aimed assessing rootstock effects on ‘Hass’ avocado. Some indirect mechanistic evidence suggests that different race rootstocks may affect postharvest anthracnose development (Willingham *et al*., 2001), alter carbohydrate accumulation (Whiley and Wolstenholme, 1990), and determine yield components, alternate bearing and nutrition (Mickelbart *et al*., 2007) on ‘Hass’ avocado. However, this study contributes new concrete evidence of direct heritable rootstock effects on new quantitative harvest and quality traits (*i*.*e*. total number of fruits, number of fruits with exportation quality, and number of fruits discarded because of low weight and thrips damage), essential for developing novel rootstock breeding schemes targeting fruit quality in the variable tropical Andes (Cortés and Wheeler, 2018).

Overwhelming rootstock effects also encourage broadening the genetic basis of current avocado rootstock breeding programs. Across Mesoamerica and northern South America, avocado trees are still cultivated in traditional orchards, backyard gardens, and as living fences. They are consumed at a regional scale, but also harbor a strong potential to improve fruit quantity and quality, besides tree adaptation, when used as rootstocks in commercial ‘Hass’ orchards (Galindo-Tovar *et al*., 2007). However, for this to occur, a better comprehension of the consequences of grafting, more concretely the rootstock-scion interaction across traits and environments, needs to be achieved, just as envisioned here.

One possible caveat of our heritability estimates refers to the number of fruits damaged by thrips. Despite it is known that rootstocks may affect the blend of biogenic volatile organic compounds emitted by ‘Hass’ (Ceballos and Rioja, 2019), and therefore influence scion pest attraction, in our study thrips’ pressure was not homogeneous across nor within orchards. In other words, different rootstocks were not equally exposed to the pest, meaning that the phenotypic vector and the relatedness matrix were fortuitously unbalanced within the ‘genetic prediction’ model. This trend was not observed for any of the other significant traits. Therefore, in order to validate the rootstock-mediated genetic-estimated heritability values obtained for the number of fruits damaged by thrips, an oncoming controlled experiment would require capturing volatiles across grafted ‘Hass’ trees, all exposed to a constant pressure by thrips.

### Relatedness reconstruction with SSR markers allow for genomic-type predictions

SSRs may not be sufficient to describe a polygenic basis but they are capable of capturing a wide spectrum of samples’ relatedness. Heterogeneity in the samples’ relatedness is essential to calibrate a ‘genetic prediction’ model when highly related or unrelated samples are not sufficiently contrasting by themselves. The molecular relationship matrix that we estimated following Lynch and Ritland (1999) and Cros *et al*. (2019) was adequately heterogeneous. In this way, our genetic prediction managed to include both family effects and Mendelian sampling terms, while simultaneously expanding the number of variables from 13 up to 240, increasing the predictive model accuracy (Zhang *et al*., 2019).

SSRs’ high mutation rate (Ellegren, 2004) and polymorphism content (Cortés *et al*., 2011; Blair *et al*., 2012) allow utilizing this type of marker for estimations of the genetic relatedness matrix, and therefore aid quantifying the additive genetic variance of quantitative traits using a ‘genetic prediction’ model. However, SSR markers will be limited when trying to assess the genomic architecture of complex traits (Hirschhorn and Daly, 2005) or when calibrating marker-based infinitesimal Genomic Selection (GS) models (Kumar *et al*., 2012; Crossa *et al*., 2017). In order to reveal the rootstock-mediated genomic architecture of scion traits, Genome-Wide Association (GWAS) models would need assuming that some rootstocks’ allelic variants are in Linkage Disequilibrium (LD) with causal variants (Hirschhorn and Daly, 2005; Morris and Borevitz, 2011; Tam *et al*., 2019) that influence the scion phenotype. Likewise, predictive rootstock breeding will have to assume that quantitative traits are regulated by infinitive low-effect additive causal variants in LD with many genetic markers (Crossa *et al*., 2017). Infrequent SSR markers, despite highly polymorphic, are definitely unlikely to be found in LD with any of these variant types (Slatkin, 2008). Hence, SNPs will needed for a deeper understanding and utilization of the rootstock-scion interaction due to their abundance and easy scoring.

## PERSPECTIVES

In order to expand our knowledge on the extent of the rootstock-scion interaction and speed up fruit tree breeding programs, further heritability estimates should be gathered on contrasting traits using multi-environment (Crossa *et al*., 2019) provenance (‘common garden’) and progeny trials with diverse panels of seedling and clonal rootstocks. The ‘genetic prediction’ model implemented here to estimate heritabilities could easily be extended to those cases at a low genotyping cost, since few SSRs markers are enough to reconstruct the genetic relatedness matrix. This model reported pervasive evidence that rootstock’s influences transcend the root phenotype and can directly impact the phenotype of the grafted scion for economically important traits. Therefore, widening the spectrum of traits under screening for rootstock-mediated heritability is essential to optimize rootstock selection and the overall genetic value of nurseries’ grafted material in the genomic era (Khan and Korban, 2012; Meneses and Orellana, 2013; Iwata *et al*., 2016).

On the other hand, rootstock-scion interaction also implies that different scions may have distinct effects on rootstock traits, such as arbuscular mycorrhizal and root hair development (Shu *et al*., 2017). Studying this type of interactions would require factorial designs in which different clonal scions are grafted ideally on clonally propagated rootstocks – *e*.*g*. via double grafting (Frolich and Platt, 1971) or micro-cloning (Ernst, 1999), or alternatively on half-sib families of seedling rootstocks. This way new scion effects can be revealed, while optimizing the rootstock-scion combination. Meanwhile, a new generation of ‘genetic prediction’ models (Crossa *et al*., 2019) may expand our understanding of how plants graft while pivoting fruit tree breeding programs. We look forward to seeing similar approaches applied on other woody perennial fruits crops as well as on orphan tropical and subtropical native trees.

Besides quantifying rootstock and scion effects using quantitative genetic approaches, a more mechanistic understanding of the consequences of grafting is desirable by applying tools from the ‘omics’ era (Barazani *et al*., 2014; Wang *et al*., 2017; Guillaumie *et al*., 2020). Genotyping-by-sequencing (Elshire *et al*., 2011) or re-sequencing (Fuentes-Pardo and Ruzzante, 2017) of each genotype, and RNAseq (Jensen *et al*., 2012; Sun, 2012; Reeksting *et al*., 2016) coupled with single-cell sequencing (Tang *et al*., 2019) across different tissues of the grafted tree, including the graft interface (Cookson *et al*., 2019), will enable understanding the genetic architecture of rootstock-mediated traits and the rootstock-scion interaction. Ultimately, these approaches may help discerning among additive and combined processes on how plant tissues and physiological processes (such as water and nutrients uptake and transport, hormone production and transport, and large-scale movement of molecules) behave during grafting.

## DATA AVAILABILITY STATEMENT

The filtered datasets and scripts are archived at Dryad Digital Repository under DOI (available upon acceptance).

## AUTHOR CONTRIBUTIONS

CAD-D, OD and AAN-A conceived the original sampling. CAD-D and OD led phenotypic data collection and root sampling. VV-Z and LP performed DNA extraction, SSR genotyping and alleles size estimation. OD and LM-B filtered and prepared input datasets. AJC, LM-B and PHR-H carried out data analyses. AJC, OD, LM-B, AAN-A and PHR-H interpreted results. AJC and PHR-H drafted a first version of this manuscript, edited by the other co-authors.

## ACKNOWLEDGEMENTS

We value M. Londoño, J.M. Cotes and M. Osorno’s insights while conceiving this project. We are also grateful to the owners and administrators of the eight avocado ‘Hass’ orchards included in this study, for allowing access to perform tree monitoring and collect root samples. Field assistants H.M Arias, J.M. Bedoya, K.Y. Calle, L.E. Cano, E. Carranza-Hernández, S.A. Guzmán, J.A. Henao, L.M. Mejía, A.M. Otálvaro, A.N. Sánchez and H.D. Yepes are recognized for collecting data and sampling roots at the eight orchards during 2015, 2016 and 2017. Special thanks for insightful discussions to G.P. Cañas-Gutiérrez, M. Casamitjana-Causa, J. Díaz-Montano, C.M. Holguín, P.E. Rodríguez-Fonseca, T. Rondón and S.M. Sepúlveda-Ortega from the SGR-funded project, and to J. Berdugo-Cely, I. Cerón-Souza and R. Yockteng from the AGROSAVIA-funded Avocado genotyping platform. Early versions of the analyses shown in this work were discussed with R. Urrea-López during the V Latin American Avocado Congress held on September 2017 in Ciudad Guzmán (Mexico), and with M. Bosacchi and S. Delphine during the 3^rd^ Global Congress on Plant Biology and Biotechnology held on March 2019 in Singapore. Some of the ideas presented in this manuscript were refined thanks to comments from A. Barrientos, J.I. Hormaza, M.F. Martínez, P. Manosalva, J. Patel and G. Wilkie during the IX World Avocado Congress held on September 2019 in Medellín (Colombia). AGROSAVIA’s Department for Research Capacity Building is thanked for supporting the participation of the last author is these events.

## SUPPLEMENTAL MATERIAL

**Table S1**. Phenotypic and genetic data of 240 ‘Hass’ avocado trees grafted on seedling rootstocks from eight orchards in the northwest of Colombia (province of Antioquia). Orchards were distributed across three agro-ecological regions, two in the dairy Northern Andean highland plateau, four in the Eastern Andean highland plateau, and two in the South West coffee region. From each orchard, 30 healthy trees from six linear blocks were chosen. Eight morphological traits were recorded in 2016, while three eco-physiological and nine harvest traits were measured from 2015 to 2017. For these last 12 traits average values across all three years are shown. Rootstocks were genotyped for 13 SSR markers (Table S2) from Sharon *et al*. (1997) and Ashworth *et al*. (2004). Alleles sizes are kept.

**Table S2**. Identity of the 13 microsatellite markers (simple sequence repeats – SSRs) used in this study to screen rootstocks from eight ‘Hass’ avocado orchards. Forward and reverse primers, sequence motif, source and summary statistics are shown. Markers were originally designed by Sharon *et al*. (1997) and Ashworth *et al*. (2004), and were prioritized according to their polymorphism information content (PIC), following Alcaraz and Hormaza (2007).

**Table S3**. Wilcoxon Rank Sum Test results comparing trait distributions across agro-ecological regions for the 20 traits surveyed at eight orchards of ‘Hass’ avocado trees grafted on seedling rootstocks. Orchards spanned three agro-ecological regions, two in the dairy Northern Andean highland plateau (heading in green), four in the Eastern Andean highland plateau (heading in red), and two in the South West coffee region (heading in orange). Traits in bold had significant heritability (*h*^*2*^) estimates (*p* < 0.05) for three different permutation strategies (of the phenotypic vector, the matrix of molecular markers and the matrix of genetic relatedness among rootstocks, Table 1).

**Table S4**. Wilcoxon Rank Sum Test results comparing trait distributions across eight orchards of ‘Hass’ avocado and examined for 20 traits within regions. Orchards spanned three agro-ecological regions, two in the dairy Northern Andean highland plateau (heading in green), four in the Eastern Andean highland plateau (heading in red), and two in the South West coffee region (heading in orange). Traits in bold had significant heritability (*h*^*2*^) estimates (*p* < 0.05) for three different permutation strategies (of the phenotypic vector, the matrix of molecular markers and the matrix of genetic relatedness among rootstocks, Table 1).

**Table S5**. Pearson correlation coefficients comparing trait distributions of 20 traits examined across eight orchards of ‘Hass’ avocado and altitude. Orchards differed in altitude (Fig. 1) and spanned three agro-ecological regions, two in the dairy Northern Andean highland plateau (heading in green), four in the Eastern Andean highland plateau (heading in red), and two in the South West coffee region (heading in orange). Traits in bold had significant heritability (*h*^*2*^) estimates (*p* < 0.05) for three different permutation strategies (of the phenotypic vector, the matrix of molecular markers and the matrix of genetic relatedness among rootstocks, Table 1).

**Table S6**. *Q*-matrix and admixture index in rootstocks from eight orchards of ‘Hass’ avocado as determined by 13 SSR markers. Estimates are derived from the STRUCTURE software (Pritchard *et al*., 2000) at *K* = 3 (Fig. 2).

**Figure S1**. Morphological first set of traits’ distributions across orchards (first column of figure panels) and agro-ecological regions (second column of figure panels) for four morphological traits (rows) – tree, trunk, rootstock and scion heights – recorded in 2016 in ‘Hass’ avocado trees grafted on seedling rootstocks at eight orchards. Orchards spanned three agro-ecological regions, two in the dairy Northern Andean highland plateau (in green), four in the Eastern Andean highland plateau (in red), and two in the South West coffee region (in orange). Orchard codes depicted in the first column of figure panels are a shorten version (last letters) of the full names shown in Fig. 1.

**Figure S2**. (Continued from Fig. S1) Morphological second set of traits’ distributions across orchards (first column of figure panels) and agro-ecological regions (second column of figure panels) for other four morphological traits (rows) – rootstock and scion perimeters, trunk perimeter at the grafting scar, and rootstock compatibility following (Webber, 1948) – recorded in 2016 in ‘Hass’ avocado trees grafted on seedling rootstocks at eight orchards. Orchards spanned three agro-ecological regions, two in the dairy Northern Andean highland plateau (in green), four in the Eastern Andean highland plateau (in red), and two in the South West coffee region (in orange). Orchard codes depicted in the first column of figure panels are a shorten version (last letters) of the full names shown in Fig. 1.

**Figure S3**. Eco-physiological traits’ distributions across orchards (first column of figure panels) and agro-ecological regions (second column of figure panels) for three eco-physiological traits (rows) – number of leaves, flowers and fruits following Salazar-García *et al*. (2013) – recorded from 2015 to 2016 in ‘Hass’ avocado trees grafted on seedling rootstocks at eight orchards. Orchards spanned three agro-ecological regions, two in the dairy Northern Andean highland plateau (in green), four in the Eastern Andean highland plateau (in red), and two in the South West coffee region (in orange). Orchard codes depicted in the first column of figure panels are a shorten version (last letters) of the full names shown in Fig. 1.

**Figure S4**. Harvest first set of traits’ distributions across orchards (first column of figure panels) and agro-ecological regions (second column of figure panels) for five harvest traits (rows) – number of fruits with exportation quality and those discarded because of mechanical damage, sun damage and damage caused by scarab beetles (*A*. *pygidialis*) or thrips (*F*. *gardeniae*) – recorded from 2015 to 2016 in ‘Hass’ avocado trees grafted on seedling rootstocks at eight orchards. Orchards spanned three agro-ecological regions, two in the dairy Northern Andean highland plateau (in green), four in the Eastern Andean highland plateau (in red), and two in the South West coffee region (in orange). Orchards codes depicted in the first column of figure panels are a shorten version (last letters) of the full names shown in Fig. 1.

**Figure S5**. (Continued from Fig. S4) Harvest second set of traits’ distributions across orchards (first column of figure panels) and agro-ecological regions (second column of figure panels) for other four harvest traits (rows) – number of fruits discarded because damage caused by *Monalonion* spp. or due to other imperfections such as low weight, early ripening or the stalk cut below the pedicel – recorded from 2015 to 2016 in ‘Hass’ avocado trees grafted on seedling rootstocks at eight orchards. Orchards spanned three agro-ecological regions, two in the dairy Northern Andean highland plateau (in green), four in the Eastern Andean highland plateau (in red), and two in the South West coffee region (in orange). Orchard codes depicted in the first column of figure panels are a shorten version (last letters) of the full names shown in Fig. 1.

**Figure S6**. Pearson correlations among eight morphological traits recorded in 2016 in ‘Hass’ avocado trees grafted on seedling rootstocks at eight orchards. Correlation estimates and 95% confidence intervals are presented above the diagonal and below diagonal cells are colored accordingly. Minimum and maximum values are shown in the corners of the cells in the diagonal.

**Figure S7**. Pearson correlations among average distributions of three eco-physiological traits recorded from 2015 to 2016 in ‘Hass’ avocado trees grafted on seedling rootstocks at eight orchards. Correlation estimates and 95% confidence intervals are presented above the diagonal and below diagonal cells are colored accordingly. Minimum and maximum values are shown in the corners of the cells in the diagonal.

**Figure S8**. Pearson correlations among average distributions of nine harvest traits recorded from 2015 to 2016 in ‘Hass’ avocado trees grafted on seedling rootstocks at eight orchards. Correlation estimates and 95% confidence intervals are presented above the diagonal and below diagonal cells are colored accordingly. Minimum and maximum values are shown in the corners of the cells in the diagonal.

**Figure S9**. Evanno’s delta *K* for the unsupervised Bayesian genetic clustering conducted in STRUCTURE and depicted in Fig. 2. *K* values ranged from *K* = 2 to *K* = 5. Transformed likelihoods of the graph model from the Evanno *et al*. (2005) are shown in the vertical axis.

**Figure S10**. Estimates of narrow-sense rootstock-mediated heritability (*h*^*2*^) in five of the 20 measured traits based on a ‘genetic prediction’ model calibrated with 13 SSRs markers genotyped in seedling rootstocks from eight ‘Hass’ avocado orchards. Depicted traits (rows) are those for which significant *p*-values (*p* < 0.05) were simultaneously obtained for three different permutation strategies (of the phenotypic vector, the matrix of molecular markers and the matrix of genetic relatedness among rootstocks, Table 1). The first column of figure panels shows the posterior distribution for the rootstock-mediated heritability (*h*^*2*^) estimates as well as their mean (dashed vertical gray line) and 95% confidence interval (continuous horizontal gray line). The second column of figure panels reflects the model fits (*R*^*2*^) expressed as the correlation between the observed trait phenotype (*y*_*i*_) and the model’s trait estimation (*βx_i_*) – equation (3). The third column of figure panels recalls the trait distribution across orchards (from Fig. 3 and Fig. S1-S5). Estimates of *h*^*2*^ and *R*^*2*^ are derived from an additive mixed linear model according to de los Campos *et al*. (2009).

**Figure S11**. Frequency distribution of pairwise estimates of Lynch and Ritland (1999)’s distance among seedling rootstocks from eight ‘Hass’ avocado orchards. Relatedness estimates were inputted in a ‘genetic prediction’ additive mixed linear model according to de los Campos *et al*. (2009) in order to compute narrow-sense rootstock-mediated heritability (*h*^*2*^) for 20 traits (Table 1, Fig. 3).

